# Expanding the RNA Virus Universe by Scalable Structure-Guided Discovery

**DOI:** 10.1101/2025.11.24.690314

**Authors:** Gaoyang Luo, Zelin Zang, Ling Yuan, Jingbo Zhou, Ao Dong, Yufei Huang, Stan Z. Li, Feng Ju

## Abstract

The discovery of RNA viruses from metatranscriptomic data remains challenging due to their extreme sequence divergence and frequent lack of conserved motifs. We present Rider, a lightweight two-stage framework that couples fast, structure-informed sequence screening with targeted structural validation. Stage 1 uses a compact 35M-parameter protein language model to prioritize RdRp-like fragments at whole-sample scale, achieving over 44× higher end-to-end screening throughput on commodity hardware. Stage 2 applies structure prediction and Foldseek-based alignment against a dedicated RdRp structure resource (∼200k ESMFold-predicted structures), providing orthogonal evidence for remote homologs. Applied to >10,000 metatranscriptomes spanning marine, freshwater, soil and host-associated microbiomes, Rider matches or outperforms leading tools (e.g., LucaProt, PalmScan) and additionally recovers divergent and truncated sequences. Multiple orthogonal indicators, including structure consistency and low DNA read mapping to corresponding contigs, support genuine RNA origin. In a human IBD cohort, Rider agrees with state-of-the-art calls for clinically relevant RNA viruses while extending discovery to divergent lineages. Rider turns structure-guided homology search into a practical, scalable pipeline for RNA virome discovery.

**Highlight:**

1. A two-stage framework enables structure-guided RNA virus discovery at sample scale, achieving up to 44-fold higher throughput on standard computing hardware.
2. The method matches or surpasses LucaProt and PalmScan across >10,000 metatranscriptomes from diverse environments, while recovering RdRp fragments missed by existing tools.
3. Structural validation using ∼200,000 ESMFold-predicted RdRp models and Foldseek alignment supports the detection of remote homologs with high confidence.
4. Orthogonal evidence, including low DNA read mapping, strand-specific expression, and ORF metrics, confirms RNA origin and reduces false positives..
5. Open-source code and an openly released RdRp structure database enable scalable, reproducible RNA virome discovery in environmental and clinical settings.

## Introduction

The rapid expansion of metatranscriptomic sequencing has greatly enhanced our ability to detect RNA viruses across a wide range of environments and host-associated microbiomes. RNA viruses are key players in microbial ecosystems and have growing relevance to human health, yet their discovery remains challenging due to their extreme sequence divergence, polyprotein organization, and frequent absence from curated reference databases. Most detection pipelines rely on sequence homology to known RNA-dependent RNA polymerase (RdRp), a hallmark gene of RNA viruses^1–3^. However, this approach struggles with novel viral families that lack canonical motifs or display fused and atypical genome structures, particularly in fragmented assemblies ^4^. Existing tools such as geNomad ^5^, DeepVirFinder ^6^, and VirSorter2 ^7^ improve sensitivity but remain limited when applied to short contigs or highly divergent proteins^8^, while popular alignment-based methods like BLAST and HMMER are further constrained due to poor sensitivity or generating false negatives ^9,10^.

Structural conservation offers a promising alternative to sequence similarity. Across diverse viral taxa, the tertiary architecture of RdRp, which is well characterized by a conserved right-hand fold, is remarkably stable despite low primary sequence similarity. Structure-based alignment methods, such as Foldseek^11^ and DALI ^12^, have thus emerged as powerful tools for detecting remote homologs that evade conventional sequence-based approaches. However, full 3D structure prediction and alignment across millions of open reading frames (ORFs) is computationally prohibitive, particularly in large-scale metatranscriptomic studies where most ORFs are incomplete, non-viral or lack clear domain boundaries. A recent state-of-the-art (SOTA) deep learning model, LucaProt ^13^, improves RNA viral detection by targeting the hall marker gene RdRp, and incorporating both sequence features and structural proxies (e.g., contact maps) into neural network architectures. LucaProt integrates Transformer-derived sequence embeddings with predicted inter-residue contact maps, which are concatenated and passed through a multilayer perception (MLP) for classification. While this design captures some aspects of tertiary structure, the generalization of whole-sample-wide protein structure from metatranscriptomic sequencing data, which typically includes millions of predicted ORFs, usually requires significant computational resources. In addition, motif-strict identification of RdRp brings high precision but may reduce performance on fragmented viral sequences. These constraints may limit the fast and sensitive identification and expansion of the RNA virome in real-world metatranscriptomic data.

Protein language models (PLMs), such as ESM (Evolutionary Scale Modeling)^14,15^, offer a promising alternative by learning latent structural and functional features directly from unannotated sequences. These models are pretrained on massive protein sequence corpora and can implicitly capture higher-order representations that correlate with secondary and tertiary structures—without requiring explicit 3D modeling or supervision. However, their application to RNA virus discovery remains constrained by several practical limitations. First, PLMs typically operate over fixed-length input windows (e.g., 1024 residues for ESM2), which can truncate long polyproteins and exclude key functional domains. Second, PLMs are not specifically optimized for viral sequence diversity or modularity and may overlook fused or rearranged domain architectures. Third, fragmented viral contigs from metatranscriptomic assemblies further disrupt domain continuity, complicating motif recognition and reducing the performance of both motif-centric and alignment-based tools. These challenges underscore the need for developing PLM-based frameworks that are robust to truncation, fragmentation, and non-canonical genome structures—hallmarks of real-world RNA virus data.

To address these challenges, we developed Rider, a two-stage framework for RNA virus discovery that integrates PLM-derived sequence embeddings (we use a compact 35M-parameter ESM encoder for efficiency) with a lightweight Transformer classifier and targeted structure-aware validation. First, to handle extreme divergence, fused domains, and fragmented contigs, Rider learns from RdRp precursors and replication-associated regions (helicases, proteases, RNA-binding), enabling motif-agnostic detection beyond canonical RdRp signatures. Second, to mitigate PLM window truncation on long polyproteins, we train on a broad length spectrum (11 to 7,334 aa) and leverage upstream replication signals so that candidates remain detectable even when RdRp motifs fall outside the input window. Third, to enable high-throughput, global-sample metatranscriptome screening without prohibitive cost, we adopt a lightweight PLM and Transformer filter to achieve >44× end-to-end speedups (including tokenization) for the screening stage alone; shortlisted hits can then undergo optional structure-aware confirmation (e.g., TM-score>0.5) ^16,17^. Meanwhile, Rider preserves SOTA-level accuracy on known RdRps (≈96%) and uniquely recovers divergent RNA viruses in real metatranscriptomes. Finally, by reducing dependence on incomplete reference databases and strict motif rules, Rider couples a fast, generalizable screening with structure-aware validation to achieve high precision, while scaling to large datasets. ^16,17^

We applied Rider to over 10,000 environmental and host-associated metatranscriptomic datasets spanning marine, freshwater, soil, plant, animal, and human microbiomes. Across these diverse ecosystems, Rider recovered substantially more RNA virus candidates—particularly in environments such as soil and freshwater (by 14.53% to 23.14 % compared with state-of-the-art methods). In the human gut virome, Rider achieved high concordance with LucaProt on clinically important taxa such as noroviruses, validating its performance against a leading state-of-the-art method. Notably, Rider also uncovered a distinct set of divergent and truncated RNA viral elements, which were often overlooked due to their atypical sequences or short lengths, highlighting the enhanced sensitivity of the method. Together, these findings demonstrate Rider’s utility as a reliable and extensible framework for RNA virome discovery in medical microbiome research. By combining PLM-derived embeddings with structure-guided validation, Rider enables scalable and comprehensive detection of deeply divergent, truncated, or embedded viral signatures across complex microbial communities.

## Methods

### Dataset construction for RNA-dependent RNA polymerase (RdRp)

RNA-dependent RNA polymerase (RdRp) is a universal hallmark gene of RNA viruses and serves as a primary target for viral discovery. However, publicly available RdRp sequences exhibit substantial variability in both sequence composition and annotation quality. Although many entries are labeled as “RdRp,” their genomic contexts vary across viral families—ranging from standalone proteins to polyproteins and truncated fragments—resulting in a heterogeneous dataset. To construct a high-confidence positive dataset, we implemented a systematic curation and filtering workflow. RdRp protein sequences were initially retrieved from the NCBI nucleotide database using the keyword “RNA-dependent RNA polymerase”. Approximately 70,000 sequences were extracted from GenBank-format files using a custom parsing script. Additionally, we also included 4803 RdRp-motif-contained sequences from the RefSeq database, where ORFs were predicted with Prodigal-gv and the clear RdRp core domains were filtered using a hidden Markov model (HMM)-based annotation^3^, with cutoff parameters: domain score > 50 and E-value < 1e−3. To reduce redundancy while preserving sequence diversity, the sequences were clustered using CD-HIT at a 95% similarity threshold, rendering 22,087 non-redundant sequences for training and testing. Note that some RdRps exist in forms of polyproteins in RNA viruses, usually followed by a series of replication-related proteins such as capsid, helicase, and protease. To enhance the generalizability of our models and address the limitation of input length (1024aa), we didn’t strictly rely on these comparatively complete proteins with the RdRp motif. Instead, we included RdRp-labeled replication-related proteins of polyproteins, as well as short partial RdRp motif fragments. These RdRp-labeled sequences were categorized into three functional groups according to their domain architecture and genomic context (Figure S2):

1. Replication-related proteins such as capsid, helicase, protease of RdRp region in polyproteins (some RdRps appear in forms of polyproteins in RNA virus);
2. Partial and complete RdRp sequences, including independent or segmented coding sequences (CDSs).

### Negative dataset construction and data partitioning

To construct a robust negative dataset, we curated protein sequences from the UniRef90 database while ensuring the complete exclusion of RdRp-related proteins. Initially, all sequences containing the terms “RdRp” or “Orthornavirae” in their annotations were removed. To further refine taxonomic filtering, we employed TaxonKit (<u>GitHub link</u>) to retrieve taxonomy identifiers from the (<u>NCBI taxdump</u>) files and excluded all sequences classified under the viral realm Orthornavirae (Taxonomy ID: 2732396).

To eliminate any residual polymerase-related sequences, we applied Palm-Scan, a domain-level classification tool capable of detecting palm-domain-containing proteins. Any sequence flagged as polymerase-like was removed, minimizing the risk of false negatives. Following preprocessing, we constructed the final training dataset with a 1:5 ratio of positive to negative sequences. For the negative training set, 110,435 non-viral non-RdRp amino acid sequences were randomly sampled from UniRef90 database (according to 1:5 ratio of positive set), and were further subdivided into 88,350 (8/10 of 110,435) for training and 4,417 (equal to positive validations, sampled from 22,087=2/10 of 110,435) for validation. Finaly, we got a training set with 17,670 positive data and 88,350 negative data and a validation set with 4,417 postive data and 4,417 negative data during training process, and a test set with 680 low similarity positive data and 200,000 negative data for comparing baseline algorithms.

### Model design, training and benchmarking

#### Model Architecture

The architecture of our model, Rider, comprises three main components: (i) a pre-trained protein language model (ESM2-35M) for contextual feature extraction, (ii) a lightweight Transformer encoder for representation refinement, and (iii) a multi-layer perceptron (MLP) for binary classification. This modular design enables sensitive detection of RdRp-like signals in protein sequences while maintaining computational efficiency.

Input sequences are first processed by the ESM2-35M model, which generates 480-dimensional contextual embeddings for each amino acid residue. To preserve the structural and evolutionary information encoded during pretraining, we freeze the weights of ESM2-35M during training. Next, the sequence of embeddings is passed to a two-layer Transformer encoder. Each layer includes multi-head self-attention (with eight heads), a feed-forward network with a hidden dimension of 960, and dropout (rate = 0.3) applied after each sub-layer. This component refines intra-sequence dependencies and captures long-range contextual relationships.

We then extract a fixed-length representation by selecting the output corresponding to the first token in the sequence, analogous to the [CLS] token in natural language models. This representation is passed to an MLP classification head, which projects the 480-dimensional vector into two logits representing the positive and negative classes. A softmax function converts these logits into class probabilities. Only the Transformer encoder and MLP layers are updated during training; the ESM2-35M backbone remains fixed. This design reduces the risk of overfitting on limited viral training data and leverages the rich priors encoded in large-scale protein pretraining.

### Model training and benchmarking

To train Rider, we fine-tuned a lightweight downstream classifier on 480-dimensional per-sequence embeddings extracted from frozen ESM2-35M protein language model. The classifier comprises a small Transformer encoder and a linear two-way output layer for binary classification (viral vs. non-viral). To address class imbalance, we applied weighted cross-entropy loss, where class weights were computed from the training data; unweighted loss was used when weights were not specified.

Training was conducted using the AdamW optimizer (learning rate = 1e−4) with a batch size of 256 per device. We employed the HuggingFace Trainer framework, with early stopping based on evaluation loss (patience = 5) and automatic selection of the best checkpoint. Logging, evaluation, and checkpointing occurred every 400 steps, and training ran for up to 1,000 epochs. During evaluation, model logits were converted to probabilities using a softmax function. A grid search over decision thresholds (0–1) was performed to identify the value that maximized F1 score, which was then used for model selection. All intermediate outputs (e.g., logits, probabilities, embeddings) and step-wise metrics were recorded for downstream analysis.

To assess performance, we benchmarked Rider against several representative models, including a standard Transformer, a CNN, a CNN-Transformer hybrid (concatenating the output embeddings of two channels), and ESM2 alone. Evaluation was performed on a balanced test set comprising 680 RdRp sequences (≤60% identity to training data) and 200,000 negatives sampled from UniRef90 using a depth-aware strategy. We further assessed performance stability through bootstrap resampling (n = 1,000). In each iteration, 200 sequences were sampled (100 positives, 100 negatives), passed through the model to extract embeddings and predictions, and used to compute ROC AUC, accuracy, F1-score, precision, and recall. ROC curves were interpolated to a common false-positive rate grid and averaged across replicates, with ±2 standard deviation bands. Final reported metrics are bootstrap means with 95% confidence intervals from the 2.5th and 97.5th percentiles of the distribution.

### Comparison of performance in identifying RNA virus

We designed a comprehensive set of benchmarking tests to evaluate the performance of Rider in comparison with baseline algorithms and SOTA tools. Our evaluation focused on two key aspects of RNA virus detection: (i) the ability to recover RNA viral sequences, and (ii) the accurate identification of RdRp sequences.

To ensure a fair comparison, we treated Rider’s structure validation module as an optional gold-standard refinement step, which can be applied to the outputs of any sequence-based method. Therefore, all benchmarking was conducted using only Rider’s sequence-based inference module. Specifically, each input sequence was embedded using ESM2-35M and passed through a lightweight Transformer-based classifier. Predictions were scored using Rider’s default threshold (≥ 0.5), and performance was measured in terms of true positive rate (TPR) and false positive rate (FPR). For FPR estimation, Rider was evaluated on 200,000 non-viral, non-RdRp proteins randomly sampled from the UniProt90 database.

#### (1) Performance on RNA virus genomes of varying lengths (NCBI RefSeq)

Metatranscriptomic assemblies are often fragmented due to the inherent complexity of transcriptomic data and limitations of current assemblers. For example, contigs assembled by MEGAHIT typically have an N50 length of approximately 500–700 bp, based on our empirical observations ^18^. Accordingly, the identification of short-fragment RNA viruses presents a significant challenge, and shorter fragments containing fewer features, such as genes. To evaluate the performance of our method on the identification of RNA virus with different length, particularly on the short length (>500bp), we tested it and compared it with other approaches (see above) using a custom-design benchmark derived from NCBI RefSeq. The benchmark includes viral genomes segmented into different length categories: 500-1000bp, 1000-2000bp, 2000-3000bp, 3000-4000bp, 4000-5000bp, >5000bp.

#### (2) Test on retrieving RNA viruses from different environments in IMG/VR database

To assess ecological generalizability, we benchmarked Rider on RNA viruses derived from nine distinct habitats in the IMG/VR (v4) database^19^. The environmental categories included: engineered, freshwater, mammals (human), mammals (non-human), marine, non-marine saline/alkaline, plants, terrestrial, and thermal hot springs. For each habitat, performance was evaluated in terms of the proportion of contigs correctly identified as viral.

#### (3) Simulated fragmented RNA virus with and without RdRp motifs

To simulate real-world conditions of fragmented RNA virus contigs, we generated Illumina-like short reads from full-length RdRp nucleotide sequences in RefSeq. These reads were assembled using MEGAHIT ^18^, and the resulting contigs were split into the same length categories as in test (1). We then used BLASTn to filter contigs containing RdRp motifs at a 95% identity threshold. This benchmark enables a controlled comparison of methods that rely on hallmark gene content, such as PalmScan and geNomad (included the marker branch of hybrid branches) ^5^.

#### (4) Identification of Known RdRp Sequences (ICTV 2021)

To evaluate sensitivity toward known RdRp sequences, we curated a dataset from the International Committee on Taxonomy of Viruses (ICTV, 2021)^20,21^. A total of 7,346 representative viral genomes were downloaded via NCBI accession numbers. Prodigal was used to predict 16,932 ORFs, from which 4,570 sequences with RdRp core domains were selected using a hidden Markov model (HMM)-based annotation ^3^. Filtering was performed with cutoff parameters: domain score > 50, domain E-value < 1e−3, and full-sequence E-value < 1e−3.

#### (5) Detection of Novel RdRp Sequences

To evaluate generalization to novel RdRp sequences, we constructed a benchmark following the novelty definitions proposed by Neri et al. ^3^. Novel RdRp sequences were categorized by taxonomic novelty at different ranks: phylum (n=117), order (n=44,595), family (n=75,604), and class (n=38,685). This test assesses each method’s ability to recover RdRp sequences that diverge significantly from existing references.

#### (6) Detection of non-RdRp RNA viral proteins

To evaluate Rider’s ability to detect RNA viral proteins beyond RdRp, we curated a dataset of non-RdRp proteins from NCBI and UniProtKB, restricted taxonomically to the RNA virus realm Orthornavirae. All sequences were manually filtered to exclude known polymerase or RdRp-associated domains. Five functional categories were selected to represent replication-related proteins: capsid proteins (n = 863), helicases (n = 417), polyproteins (n = 2,049), viral proteases (n = 521), and other annotated proteins (n = 3,649). These categories reflect functional diversity beyond canonical motifs and test Rider’s broader generalization ability.

### Runtime benchmarking of deep learning-based protein annotation methods

To systematically compare the runtime performance of Rider and LucaProt ^13^, we constructed a controlled benchmark dataset and evaluated both methods under standardized conditions. We curated a set of protein sequences from the UniRef90 database, explicitly removing any entries annotated as RdRps to ensure a negative dataset. Sequences were filtered to retain only those with a minimum length of 1024 amino acids. From this filtered pool, we randomly selected the 1,000 sequences and constructed nine subsets of increasing size (100 to 900 sequences, in increments of 100). Each subset was named accordingly (e.g., sample_100, sample_200, etc.) and used for runtime evaluation. For each subset, we evaluated the inference time of Rider and LucaProt, excluding their downstream structure validation modules to ensure a fair comparison of the core deep learning-based annotation components. All input sequences were truncated or padded to a uniform length of 1024 amino acids. Both methods were executed on the same hardware environment, using a single NVIDIA A100 GPU with 80 GB of memory. GPU memory usage was controlled to remain below 12 GB during execution to ensure comparable resource constraints (for Rider with the parameter of input batch size 64). The runtime of Rider was recorded from its internal logging system, including timing information for each processing step and the total execution time. For LucaProt, runtime was measured via system-level wall-clock timing, capturing the start and end times of the inference process. Each method was run once per dataset subset. The total runtime (in seconds) was recorded for comparative analysis. Rider demonstrated consistently faster inference, achieving an average of ∼44-fold speedup over LucaProt across all dataset sizes.

### Construction of RNA virus protein structure library

To support structural validation of putative RNA virus proteins, we constructed a large-scale structure library comprising diverse RNA virus-encoded proteins, namely the Rider RdRp Structure Database. Protein sequences were sourced from the four data sources: NCBI ^22^, Neri et al.^3^, Zayed et al.^23^, Edgar et al.^2^. However, upon inspection, this dataset was found to include a broad range of viral proteins beyond canonical RdRps ^22^, including helicases, proteases, and capsid proteins. Rather than filtering these out, we retained the full diversity of viral replication-associated proteins to enable a more comprehensive structural comparison, as these signals may appears as precursors of RdRps in polyproteins. We used ESMFold v1, a high-accuracy deep learning model for protein structure prediction, to generate 3D models for each sequence. Following standard protocol, only the first 1024 amino acids of each sequence were used as input, with longer proteins truncated and shorter ones padded. The resulting structures were stored in PDB format and compiled into a reference structure library. The final database contains 229,721 predicted structures representing a wide array of RNA virus proteins and was used for structural homology estimation and validation of Rider’s predictions.

### Validation of candidate proteins using fast structure alignment

The challenges associated with interpreting the output of neural network predictions, often referred to as the "black box" problem, make it difficult to generate convincing validation results. To address the interpretability challenges of neural network-based predictions, we implemented a structure-based validation strategy using Foldseek^11^, a rapid structural alignment tool. Specifically, we adopted the Probability Score (Prob-score), a probabilistic metric introduced by Foldseek (TM-Score)^11^, to assess the likelihood that a structural match represents a true homologous relationship. For each Rider-predicted candidate, the amino acid sequence was first converted into a 3D structure using ESMFold v1, truncated to 1024 amino acids if necessary. The predicted structure was then aligned against our curated RNA virus protein structure library (see above) using Foldseek. Prob-score was calculated as the posterior probability of a structure match being a true positive (TP) given the observed bit score, defined as:

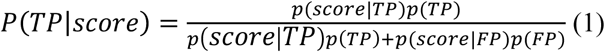

where TP and FP represent true and false positives, respectively, and the probabilities were estimated from empirical score distributions.

To benchmark this approach, we constructed a positive reference set comprising 229,721 known or putative RdRp protein structures from the source data for constructing NeoRdRp2 database^22^, and a negative reference set of 200,000 non-RdRp proteins randomly sampled from UniRef90. All structures were predicted using ESMFold under consistent parameters. First, we conducted an all-vs-all Foldseek alignment within the positive set. Matches between known RdRps yielded high structural similarity scores (typically >100), confirming strong internal homology. Next, we aligned the negative set against the positive reference. These comparisons yielded significantly lower scores, with very few exceeding benchmark thresholds (e.g., score > 50 or TM-Score > 0.5, based on Foldseek’s SCOPe40 evaluation), indicating high specificity. We uniformly applied a Prob-score threshold of 50 as a moderate but reliable cutoff for confident identification of structurally validated candidates.

### Validation of Foldseek-based structural scoring

We benchmarked the structural validation workflow using our curated Rider RdRp structure database, which contains 229,721 known or putative RdRp proteins (modeled with ESMFold; inputs truncated to 1,024 amino acids where needed). For specificity assessment, we additionally assembled a negative set of 200,000 non-viral, non-RdRp proteins sampled from UniRef90 and modeled all sequences with ESMFold under identical settings. Each query structure (positive or negative) was aligned to the Rider database with Foldseek (in ‘easy-search’ mode with ‘--alignment-type 1’), and structural evidence was summarized as the mean of the Top-K probability-like scores (Prob-scores) per query (Ave_Prob_score = TopK_hit_Prob_score/K), with K = 10 by default. The default K requires concordant support from multiple high-ranking neighbors and mitigates single-hit outliers and mixed-domain or truncated queries; when coverage is limited, the same aggregation can be applied with smaller K (1–9), while K = 10 remains the recommended setting to demonstrate structural conservation (i.e., ≥10 similar structures present in Rider). The decision threshold was set at Prob-score = 50, consistent with prior Foldseek calibration and confirmed by our benchmarking (based on Foldseek’s SCOPe40 evaluation ^11^). The score distributions were distinctly bimodal (Figure S1): negatives concentrated at low values near 20, whereas positives peaked at high values (typically >80, approaching 100). This pattern is consistent with strong internal homology among RdRp structures.

### DNA read-based validation of metatranscriptomic contigs

To evaluate the genomic validity of Rider-predicted RNA virus contigs, especially novel ones^3,13^, metagenomic DNA reads were mapped to contigs assembled from metatranscriptomic data. Paired metagenomic and metatranscriptomic datasets were obtained from four sources:

- Activated sludge samples (n = 11): collected from ten municipal wastewater treatment plants.
- Soil samples (n = 18): generated in-house using Illumina sequencing.
- Human gut microbiome samples (n = 39): retrieved from the IBD cohort of the Human Microbiome Project.
- China environmental metatranscriptomic dataset (n = 50): derived from the LucaProt validation dataset ^13^.

For each sample, metatranscriptomic reads were quality-filtered using fastp (v0.23.1) ^24^ and assembled into contigs using MEGAHIT ^18^. To quantify the genomic support for each contig, we calculated the DNA mapping rate, defined as the proportion of nucleotide positions covered by at least one DNA read. Metagenomic DNA reads were aligned to metatranscriptomic contigs using Bowtie2 (v2.3.5.1)^25^ with default parameters. Coverage statistics, including the DNA mapping rate, were computed using the ‘pileup.sh’ utility from BBMap (v39.01)^26^, which reports the percentage of positions with non-zero read depth (i.e., coverage ≥ 1) relative to the total contig length. The DNA mapping rate serves as an orthogonal indicator of transcript validity—higher coverage implies that the RNA contig is likely derived from a DNA-encoded viral genome, rather than from assembly artifacts or transient RNA molecules.

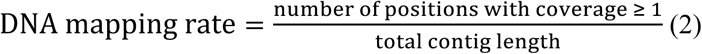

Mapping coverage was defined as the percentage of contig bases covered by at least one DNA read.

Coverage statistics and RPKM (Reads Per Kilobase per Million mapped reads) values were computed using bbmap ^27^, following the standard formulation:

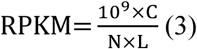

Where C: number of reads mapped to a contig with Rider hit ORFs. This represents how many sequencing reads align to the gene of interest. N: total number of mapped reads in the experiment. This includes all reads mapped to all genes or transcripts and is used to normalize for sequencing depth. L: length of the gene or transcript in base pairs (bp). This accounts for gene length bias, as longer genes are more likely to accumulate more reads.

For human gut samples, additional quality control was applied to remove interspersed repeats and low-complexity sequences using RepeatMasker (v4.1.8) ^28^ with the Dfam 3.9 database (dfam39_full.7) ^29^.

To investigate the relationship between Rider’s prediction confidence and DNA support, contigs were stratified into five confidence bins based on Rider-assigned scores: 50–60, 60–70, 70–80, 80–90, and 90–100. For each bin, DNA mapping rates (mapping sites / whole length) were analyzed using kernel density estimation (KDE) implemented in Seaborn (v0.11.2, Python). This enabled visualization of the correlation between prediction confidence and the likelihood of corresponding DNA reads, serving as an orthogonal validation of the predicted viral sequences.

### Structural-based taxonomic annotation of Rider-predicted viral contigs

To assign taxonomic labels to Rider-predicted viral ORFs for community-level profiling (Figure 5j), we constructed a custom reference database of RNA viral proteins enriched for both canonical (e.g., RdRp) and non-canonical viral genes. The database was assembled from NCBI Virus protein records, including sequences from complete RNA virus genomes across diverse taxa. All proteins were clustered at 90% amino acid similarity to reduce redundancy. We used Foldseek (v4.0) in ‘easy-search’ mode with ‘--alignment-type 1’ (local alignment) to perform pairwise structural comparisons between Rider-predicted ORFs and the reference viral protein database. For each ORF, the top-scoring hit (highest bit-score) with a minimum bit-score ≥ 50 was retained. The corresponding NCBI taxonomy ID from the best hit was assigned to the query ORF. Given the absence of a standardized taxonomy framework for many RNA viruses, especially in metatranscriptomic datasets, we adopted a family-level annotation strategy to ensure interpretability and consistency across analyses. Only ORFs with confident structural matches and low and moderate DNA mapping rates (typically <90%) were retained for downstream taxonomic profiling.

### Implementation of environmental and human metatranscriptomes

To evaluate the real-world performance of Rider in RNA virus discovery, we applied it to large-scale environmental and human virome datasets and compared its predictions to two state-of-the-art (SOTA) motif-based methods: LucaProt ^13^ and PalmScan ^1,2^. This analysis was conducted across two application domains: (1) environmental metatranscriptomic data and (2) human gut virome data from an inflammatory bowel disease (IBD) cohort.

#### Environmental metatranscriptomic data

Environmental RNA virome detection was conducted on a comprehensive dataset comprising over 10,000 metatranscriptomic samples collected across 45 ecosystems worldwide, including soil, marine, freshwater, and wastewater environments ^13^. These samples were obtained from the LucaProt benchmarking release and represent one of the most extensive environmental metatranscriptomic resources to date. ^13^ For each sample, raw metatranscriptomic reads were subjected to quality control using fastp (v0.23.1), followed by de novo assembly with MEGAHIT. Open reading frames (ORFs) were predicted using Prodigal (in metagenomic mode), and sequences were filtered to retain only those between 100 and 2000 amino acids ^13^. To reduce redundancy, predicted viral-like ORFs were clustered at 90% amino acid similarity using MMseqs2, and representative centroid sequences with a cluster with more than 4 items (≥4) were used for prediction and comparison. Each ORF was then scored by Rider (Prob), LucaProt (--threshold >0.5), and PalmScan independently. Rider prediction scores were used to stratify viral candidates into five confidence intervals (50–60, 60–70, 70–80, 80–90, and 90–100). To validate Rider’s capacity to detect canonical RdRp sequences, we benchmarked its performance against a previously published environmental RdRp dataset comprising 161,979 known or putative RdRp-containing sequences. Rider’s recall rate was calculated as the proportion of known RdRps successfully identified within this dataset.

#### Human gut virome analysis in IBD cohort

To investigate the applicability of Rider in a disease-relevant context, we applied it to a well-characterized IBD cohort, including samples from Crohn’s disease (CD, n=92), ulcerative colitis (UC, n=21), and non-IBD (healthy controls, n=11)^30^. Paired metagenomic and metatranscriptomic datasets were obtained for each subject. Metatranscriptomic reads were processed using the same pipeline as for the environmental data, including quality filtering with fastp, assembly with MEGAHIT, and ORF prediction via Prodigal. For human gut samples, additional filtering was performed using RepeatMasker (v4.1.8) with the Dfam 3.9 database to remove interspersed repeats and low-complexity sequences. Rider was used to score all predicted ORFs, and candidate viral sequences were filtered by length to isolate short proteins (100–200 amino acids), which are often missed by conventional methods. These short candidates were subjected to structure-based validation using ESMFold and Foldseek, with Prob-score used to assess structural similarity to known viral proteins. To assess the genomic plausibility of predicted contigs, paired metagenomic DNA reads were mapped to assembled metatranscriptomic contigs using Bowtie2. DNA mapping coverage and RPKM values were calculated using bbmap. Contigs with high DNA mapping rates (>90%) were flagged as potential endogenous or DNA virus-like elements, while those with low or no DNA support were considered likely to originate from RNA viruses.

### Protein sequence similarity and structure visualization

Global pairwise sequence similarity was computed with the Needleman–Wunsch algorithm (Biopython, Bio.pairwise2; default gap penalties unless noted). Protein structures were saved in PDB format and visualized in PyMOL (Schrödinger, LLC).

### Statistical analysis

All statistical analyses were performed using Python (v3.10.18) and R (v4.2.1) with standard scientific computing libraries. For comparisons of viral abundance and diversity metrics between sample groups, non-parametric tests were used unless otherwise specified. Alpha diversity (within-sample diversity) was measured using the Shannon index, calculated via the vegan package in R. Beta diversity (between-sample dissimilarity) was assessed using Bray–Curtis dissimilarity, followed by principal coordinate analysis (PCoA) for dimensionality reduction and visualization. Group differences in diversity and abundance were evaluated using the Mann–Whitney U test for pairwise comparisons and Kruskal–Wallis H test for multiple groups. Statistical significance was defined as p < 0.05, unless otherwise stated. All p-values were adjusted for multiple testing using the Benjamini–Hochberg false discovery rate (FDR) method where appropriate. To assess the correlation between prediction confidence scores and DNA mapping support, kernel density estimation (KDE) was performed using Seaborn (v0.11.2) in Python. KDE plots were stratified by Rider-assigned confidence intervals to visualize the relationship between predicted viral likelihood and supporting DNA evidence. For structural validation, Prob-score distributions were compared across groups or validation sources. In particular, mean Prob-scores were used as quantitative indicators of structural homology between Rider-predicted candidates and known viral proteins. Structural alignment statistics were visualized using matplotlib and seaborn. All plots and statistical outputs were generated in reproducible scripts and are available upon request or via the project’s GitHub repository (see Data and Code Availability section).

### Code availability

Rdier is open-sourced under the MIT license. The code repository is available from GitHub (https://github.com/emblab-westlake/Rider).

## Results

### Rider: A deep representation model for diverse RNA virus discovery

Rider is a lightweight deep learning framework designed for RNA virus identification in metatranscriptomic data. At its core, Rider leverages ESM-2, a pretrained large protein language model with 35 million parameters, to extract protein embeddings that encode both local motifs and global structural features. These embeddings are then processed by a Transformer encoder, which captures long-range dependencies and contextual information. To assess Rider’s classification performance, we benchmarked it against a range of baseline models and state-of-the-art algorithms. On a balanced test set containing 680 non-redundant RNA virus proteins and 200,000 non-viral proteins, Rider achieved an area under the ROC curve (AUC) of 0.989 (Figure 2a), outperforming all tested baselines, including CNN and Transformer-based models. Rider also consistently delivered higher accuracy, F1-score, recall, and precision (Figure 2b), indicating robust classification across diverse sequence backgrounds. To enable fast and scalable inference, Rider uses a compact architecture that can process over 500,000 sequences in 0.4 seconds on an NVIDIA A100 GPU, approximately 44 times faster than SOTA on large-scale datasets (Figure 2c), making it suitable for large-scale metatranscriptomic screening. We visualized Rider’s latent representations using t-SNE, UMAP, and PCoA, consistently revealing clear separation between viral and non-viral proteins (Figure 2f). This suggests that Rider learns generalizable sequence features beyond canonical RdRp motifs.

Secondly, we benchmarked on Rider’s ability to recover RNA viral fragments. For the NCBI RNA virus benchmark, we split original nucleotide sequences into length groups (base pairs or bp) to satisfy different input requirements (see Figure 2 Note). Unlike alignment-based or motif-driven tools, Rider is optimized to detect short or fragmented sequences that are common in real-world metatranscriptomic assemblies and often missed by traditional methods (Figure 2e). Note that shorter fragments may lack several motifs (for example Virsorter2 is feature-number-dependent and LucaProt is RdRp-motif-specific-dependent). With length increased, each method was comparable in performance (>5000bp). We further validated Rider’s ability to recover RNA viral sequences from nine ecological biomes curated from the IMG/VR database, including marine, freshwater, terrestrial, and host-associated environments. Across all biome categories, Rider achieved the highest true positive rates (TPRs) (Figure 2d), while maintaining high precision (97%) (Figure 2b), confirming its adaptability to diverse viral sequence compositions occurred in environmental microbiomes.

To further benchmark Rider’s performance on canonical RNA virus markers, we evaluated its ability to detect RdRp sequences curated from the ICTV 2021 reference. Rider correctly recalled 4,338 out of 4,570 annotated RdRp proteins (95.0% recall), closely matching LucaProt’s 98.1% recall (Supplementary Excel Table S1). Despite being a lightweight model trained on a broader virus classification task, Rider retains strong sensitivity on this well-established target. We also assessed generalization to novel RdRp sequences across increasing levels of taxonomic divergence, following the protocol proposed by Neri et al. ^3^ Rider achieved an average recall of 84.7%, corresponding to 93.4% of LucaProt’s performance across phylum, order, family, and class-level novelty groups (Supplementary Excel Table S2). These results confirm that Rider maintains robust detection capabilities even for highly divergent RNA viruses, while offering substantial improvements in speed and scalability over existing methods.

### Rider identifies remote homologs RdRp via structure-aware validation

To evaluate Rider’s utility in real-world virus discovery, we applied it to a published metatranscriptomic dataset of activated sludge, a microbial-rich environment in wastewater treatment systems. This engineered ecosystem is known to harbor diverse and under-studied RNA viruses, making it a challenging but informative testbed. The dataset comprised over 120 million paired-end reads from complex microbial communities. Within this dataset, Rider identified a highly divergent picorna-like RNA virus candidate (contig AD_AS_287816; Figure 3a) that would have, otherwise, escaped detection by existing tools. Despite sharing only 40.6% sequence similarity with the closest known reference, Rider confidently classified the predicted RdRp as viral. Supporting this classification, structural alignment with a known RdRp (UVM79539) revealed 98.0% structural similarity, with conserved catalytic motifs A–C clearly resolved in three dimensions. This case exemplifies Rider’s capacity to detect deeply divergent, functionally conserved proteins potentially missed by traditional alignment- or motif-based tools.

A key advantage of Rider lies in its superior ability to identify remote homologs using representations that capture both structural and functional features, rather than relying solely on primary sequence similarity. Many RdRps from positive-sense RNA viruses are embedded within larger polyproteins and exhibit high sequence variability, with lengths ranging from ∼460 to ∼1930 amino acids depending on the viral family. In such cases, the N- and C-terminal boundaries of catalytic domains are often ill-defined, making motif-based recognition less reliable. Rider overcomes these limitations by leveraging distributed semantic and structural signals across full-length or fragmented sequences, enabling robust detection of functionally relevant but sequence-divergent viral proteins.

To support the structural filtering step in Rider’s pipeline (see Figure 1, Step 3), we constructed a benchmark of positive and negative protein structures and evaluated structural similarity using Foldseek. As shown in Figure S1, the probability score distributions for known RdRp and non-RdRp proteins exhibit clear separation. A similarity cut-off of 50 TM-score for structure similarity was chosen to distinguish true positives from background noise. While some false positives remain above this threshold, they are largely filtered out by Rider’s upstream Transformer-based classifier, preventing them from progressing to the structure validation stage. This layered strategy improves both specificity and computational efficiency in high-throughput discovery scenarios of RNA viruses from large-scale (meta)genomics data.

**Fig. 1.**
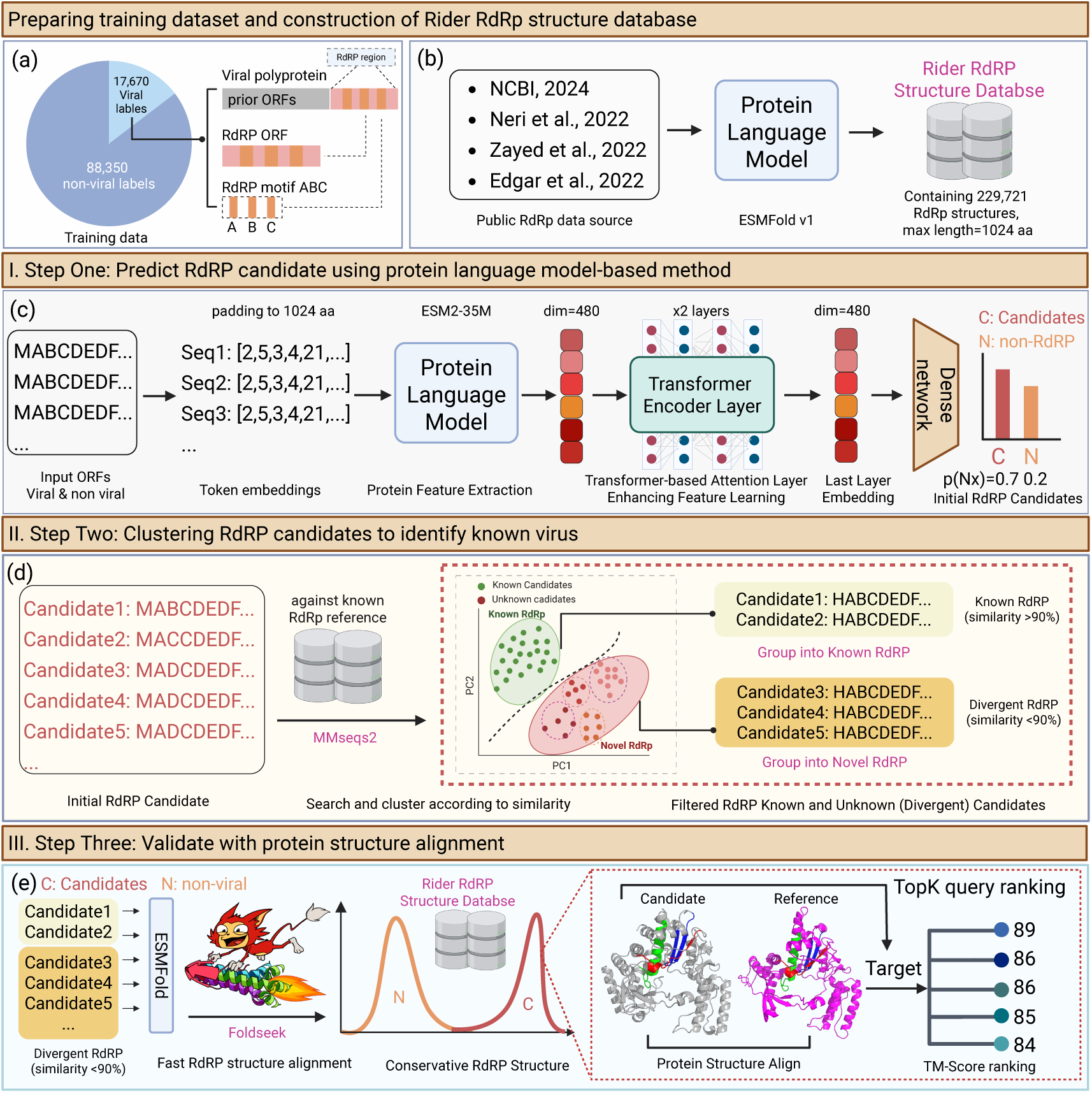
Overview of the Rider workflow for fast and sensitive RNA viruses discovery. This figure illustrates the Rider’ multi-step workflow for identifying and validating potential novel RNA-dependent RNA polymerase (RdRp) proteins from metatranscriptomic data. **a**, Training and validation date settings. **b**, Construction of Rider RdRp structure database, sourced from public datasets. (NCBI ^22^, Neri et al.^3^, Zayed et al.^23^, Edgar et al.^2^). **c**, Step one: Open reading frames (ORFs) are tokenized and embedded using a protein language model. A Transformer layer refines the feature representation, and a dense network outputs probability scores to classify sequences as viral and non-viral. **d**, Step two: High-confidence candidates are filtered via sequence similarity against the known RdRp reference. Sequences with <90% similarity to known RdRps are retained as divergent candidates. **e**, Step three: All candidates were used to generate 3D structure, then structural filtering is performed using Foldseek to compare predicted structures against Rider RdRp structure database. TopK-ranking (top k possible targets according to the alignment probability score, or TM-Score) candidates are selected based on structural similarity for further validation.

**Fig. 2.**
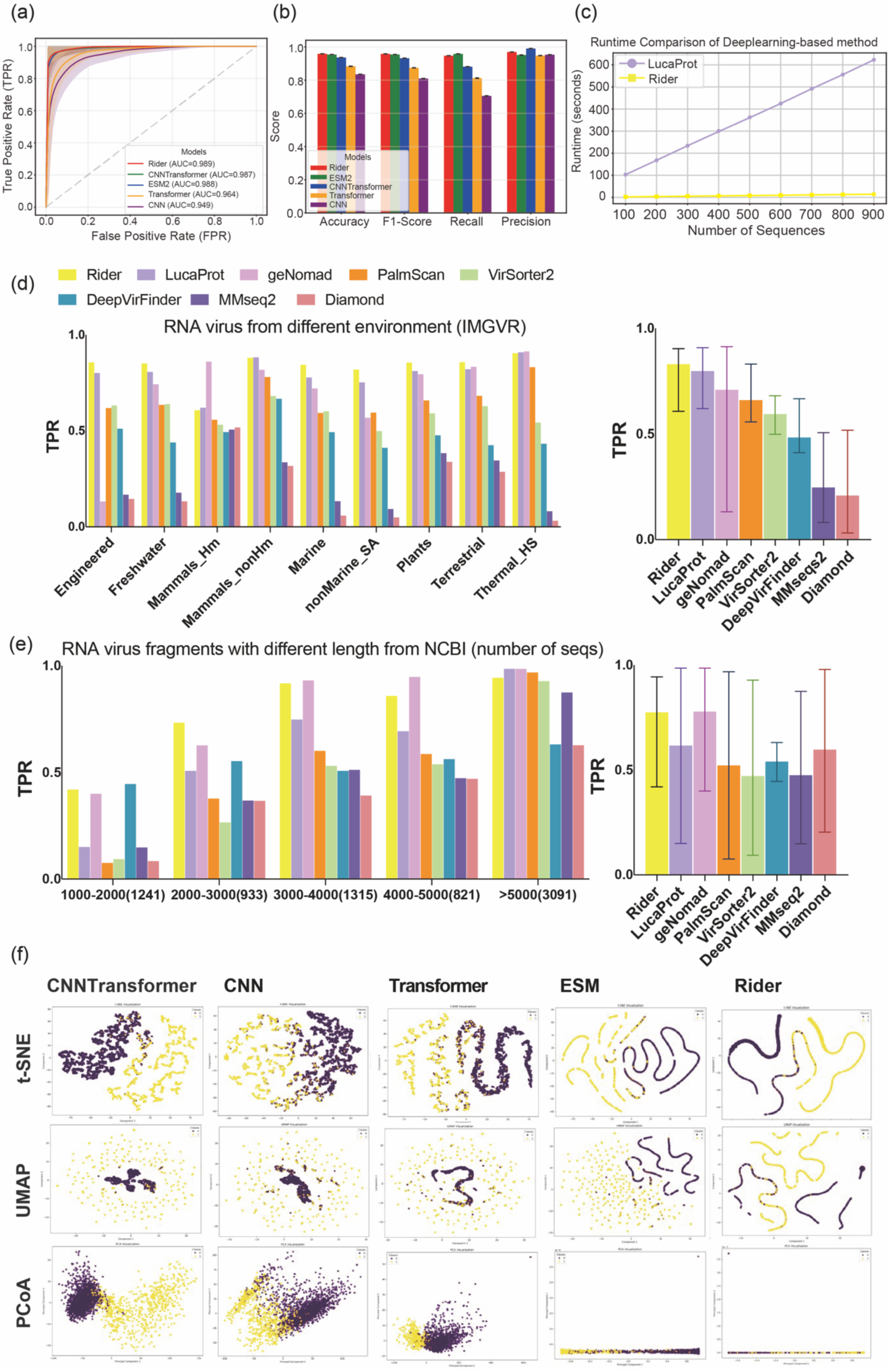
Benchmarking the Rider family against state-of-the-art methods. **a**, ROC curve comparison for different models, showing true positive rate (TPR) versus false positive rate (FPR), with the Rider algorithm achieving competitive performance. **b**, Model performance metrics including Accuracy, F1-Score, Recall, and Precision, with error bars indicating variability. **c**, Runtime comparison between two deep learning-based methods, Rider (with 34M parameters including PLM part) and LucaProt (with 24M parameters), with truncation length set to 1024 amino acids and comparable GPU memory usage (∼12GB). LucaProt consists solely of a deep neural network, whereas Rider is a full pipeline that includes an optional downstream structure-validation module. For a like-for-like assessment of core inference speed, we benchmark only the deep learning component of Rider and omit its optional module. This controls pipeline heterogeneity and isolates the contribution of the neural architectures under identical I/O and batching. Rider demonstrates significantly higher efficiency, running approximately 44 times faster than LucaProt. **d**, Benchmarking results on the IMG/VR v4 database, comparing the true positive rate (TPR) across different viral taxonomy groups for Rider and other SOTA methods. **e**, Benchmarking results on RdRp sequences from the RefSeq database with various length ranges (nucleotide base pairs, see *Notes.), evaluating TPR across different sequence length ranges and taxonomy groups. Error bars represent variability across datasets. **f**, Visualization of model representations using different dimensionality reduction techniques. The columns correspond to different models (Rider-CNN, Rider-Transformer, Rider-CNN/Transformer, Rider-ESM2, and Rider), while the rows represent different dimensionality reduction methods (t-SNE, UMAP, and PCA). The color gradient indicates variations in feature space distributions, highlighting differences in how each model captures underlying data structures. *Notes: Recall were calculated if the nucleotide sequence or any protein predicted within the sequences were identified. geNomad^5^, DeepVirFinder ^6^, and VirSorter2 ^7^ use nucleotide as input, the rest of methods use amino acid as input. All amino acids will return to nucleotide index; X axis of (e) denotes length range (number of fragments within each group).

### Learning viral signatures beyond RdRp: structural and functional generalization

To evaluate whether Rider can recognize RNA virus proteins beyond canonical RdRp domains, we tested three classes of non-RdRp proteins: capsid, helicase, protease, and polyprotein. Note that helicases and proteases, and in some cases capsid proteins, are frequently located upstream of the RdRp domain within large viral polyproteins. Although Rider was trained on RdRp-labeled sequences, this co-occurrence likely exposed it to upstream replication-associated signals during training. Rider achieved high recall across these categories (polyproteins 100%, helicases 57.1%, proteases 68.0%, capsid proteins 67.3%; Figure S4), indicating that the model has learned generalizable viral replication features. These distributed signals, including helicase, protease, RNA-binding motifs and membrane anchors features and capsid structural signals, enable Rider to detect viral proteins even when canonical RdRp motifs are absent from the input window, indicating a practical advantage for identifying fragmented metatranscriptomic assemblies and contigs lacking coverage of RdRp region.

This finding is consistent with the genomic organization of many RNA viruses, which often encode their replication machinery as large polyproteins subsequently processed into functional subunits. For example, the RdRp complex in MERS-CoV resides within a polyprotein that requires proteolytic cleavage for activation ^31^. Similarly, viral RdRps are frequently fused with helicases, methyltransferases, or proteases, forming modular replication units that vary across viral families and are often under-annotated ^32^. Among these, helicases—such as nsp13 in coronaviruses—are among the most conserved replicative enzymes and form essential components of the replication complex in nidoviruses ^33^. Rider’s ability to recover these proteins, including partially annotated or polyprotein-embedded domains, suggesting that it may capture latent structural and functional signals associated with viral replication.

Rider does not require the presence of canonical RdRp motifs within the input window to flag a sequence as viral. For instance, the Yak enterovirus polyprotein (YP_009243645.1) contains an RdRp domain (residues in 1714–2166 region) as well as upstream helicase, capsid, and protease regions. Rider recovered this entire region from those precursors, while LucaProt only detected the protein when canonical RdRp motifs were present. Similarly, Sesbania mosaic virus polyprotein P2a (NP_066392.4), which lacks an RdRp domain but encodes a protease, VPg, ATPase (P10), and RNA-binding proteins that are essential for genome replication and movement ^34^, was confidently predicted by Rider but missed by LucaProt. ^35^ Importantly, this behavior generalizes to large nidoviral polyproteins, for example, in Coronaviridae in which the RdRp is encoded as non-structural protein 12 (nsp12) within ORF1ab and can lie well downstream of ORF1a. To systematically evaluate this behavior, we analyzed 109 annotated ORF1ab or ORF1a polyprotein sequences from Coronaviridae (Table S3). Among these, 12 sequences corresponded to ORF1a fragments lacking the canonical RdRp region (nsp12). All 12 were confidently predicted as viral by Rider, whereas LucaProt failed to detect all cases. For the remaining 97 ORF1ab sequences, LucaProt failed to detect 25 cases, likely due to the RdRp domain lying beyond its 4000-residue input limit. In contrast, Rider correctly identified 104 sequences as positive, demonstrating its ability to generalize from upstream replication-associated signals even when the canonical RdRp domain is absent or inaccessible. For instance, both the full ORF1ab replicase of Magpie-robin coronavirus (HKU18) and its ORF1a fragment (residues 1–3614) are retrieved by Rider, whereas LucaProt fails on the ORF1a fragment because the RdRp core (nsp12) resides at residues ∼3609–4536 and falls outside LucaProt’s detection window. These results reinforce Rider’s robustness in recovering long, modular viral polyproteins that exceed the receptive field of motif-centric models.

Structural comparisons further support Rider’s detection of non-canonical viral proteins. In a contig from activated sludge metatranscriptome assemblies (AD_AS_287816), Rider detected both a canonical RdRp and a second protein structurally similar (77%) to a known picorna-like capsid protein (YP_009337257.1), despite only 17.3% sequence similarity (Figure 3b). Extending analysis to intermediate-scoring proteins (Rider score 50–70), we identified structural matches to capsids from *Apis Dicistrovirus*, Antarctic Picorna-like virus 1, and *Squash mosaic virus* (Figure S3), with Foldseek homology probabilities of 53–68%. Complementarily, 3.6% (725/20,000) of training sequences showed high similarity (>60%) to annotated non-RdRp proteins (polyproteins n=373; helicases n=143; proteases n=112; and capsids n=97), reflecting the modular, multifunctional nature of many RNA virus polyproteins and exposing the model during training to a diversity of replication-associated features. Together, these structural and dataset analyses support the conclusion that Rider is sensitive to distributed replication-related signals that extend beyond canonical RdRp motifs.

**Fig. 3.**
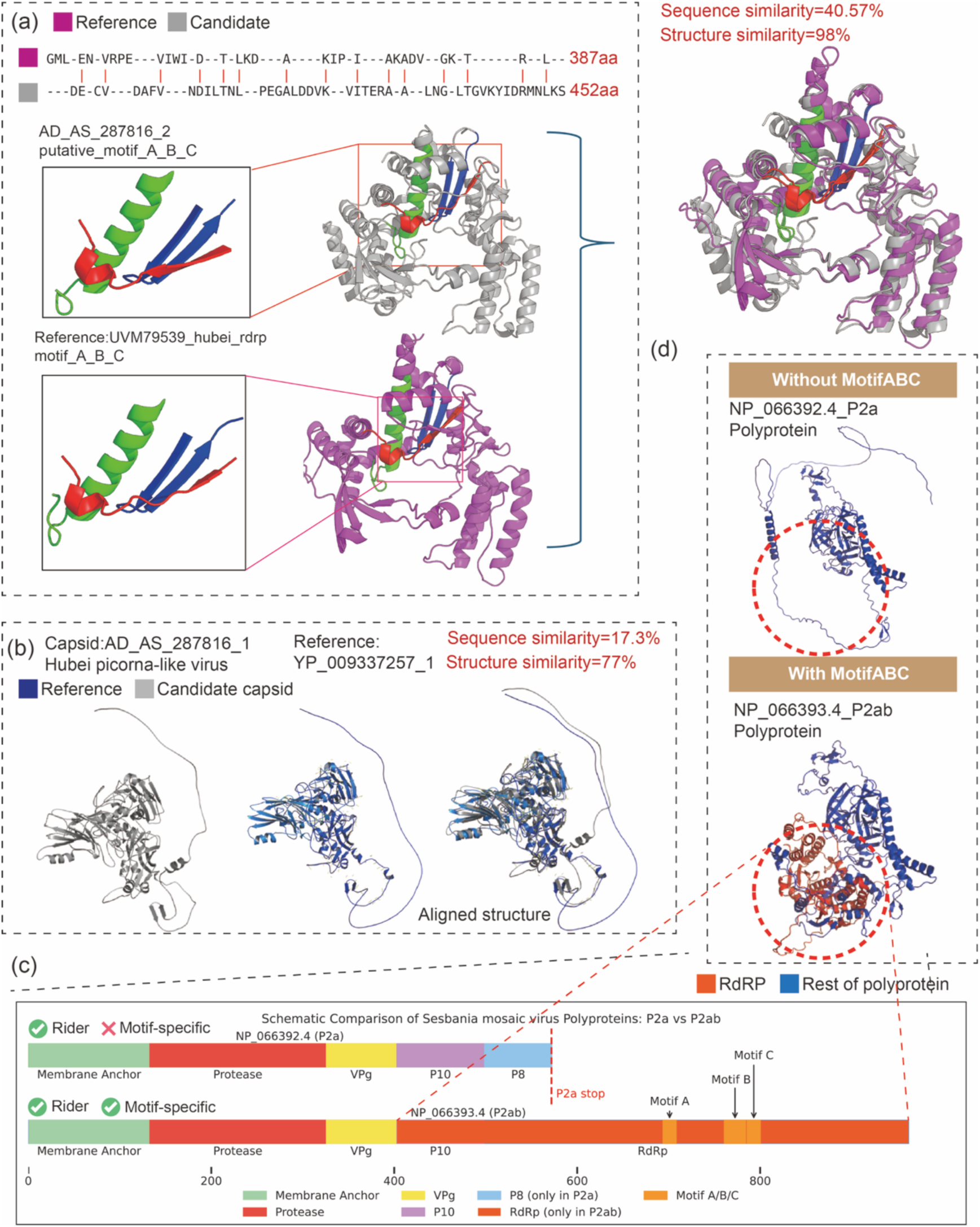
Structural validation and domain-aware analysis of Rider-predicted RNA virus proteins. **a,** Structural alignment between a Rider-predicted candidate (AD_AS_287816_2) and a known RdRp from UVAT05730_hubei_rdrp reveals high structural similarity (77%) despite low sequence similarity (40.57%). Conserved motifs A–C are accurately recovered in both structures (highlighted in green), demonstrating Rider’s ability to identify deep homologs with divergent sequences. **b**, Structural comparison of a Rider-predicted capsid protein (AD_AS_287816_1) from a Hubei picorna-like virus with a known reference (YP_009337257_1) shows strong alignment, confirming that non-RdRp RNA virus proteins detected by Rider have strong conservative structure, while this protein may have low sequence similarity (17.3%). **c**, Domain-level comparison between two Sentinel mosaic virus polyproteins (P2a vs P2ab). Rider correctly identifies both RdRp and adjacent replication-associated regions, while motif-specific method (here LucaProt) detects only canonical motif-containing sequences. The motif A–C region is absent in one polyprotein (P2a), highlighting cases where Rider captures broader replication signals missed by motif-based tools. **d**, Structural validation of two polyproteins using ESMFold predictions. The polyprotein lacking MotifABC (NP_069892.4_P2a) shows no detectable RdRp fold, while the one containing MotifABC (NP_069893.4_P2ab) aligns well with known RdRp structures. This further supports the utility of structural features in validating partial or divergent replication proteins.

### DNA read coverage supports Rider predictions

To evaluate whether Rider-predicted RNA virus candidates correspond to genuine genomic entities, we performed orthogonal validation using DNA read coverage from 80 paired metagenomic and metatranscriptomic datasets of environmental samples. Predicted candidates with over 50 Prob score after structure validation were included in this section. This multi-omics approach enabled us to assess whether high-confidence RNA-based predictions are supported by the corresponding underlying DNA signals, as recommended by Xin et al.^36^. Across all paired datasets, a consistent trend emerged: contigs with higher Prob-scores tended to exhibit lower DNA mapping rates (Figure 4e-h). For instance, contigs scoring between 90–100 showed the lowest DNA coverage (<5%). The distribution of DNA mapping rates was right-skewed, with a mean of 3.5%, reflecting the expected scarcity of DNA evidence for RNA-dominant or fragmented contigs. This pattern was consistently observed across diverse environmental samples, including soil and wastewater, reinforcing the biological relevance and reliability of Rider predictions in real-world metatranscriptome contexts.

**Fig. 4.**
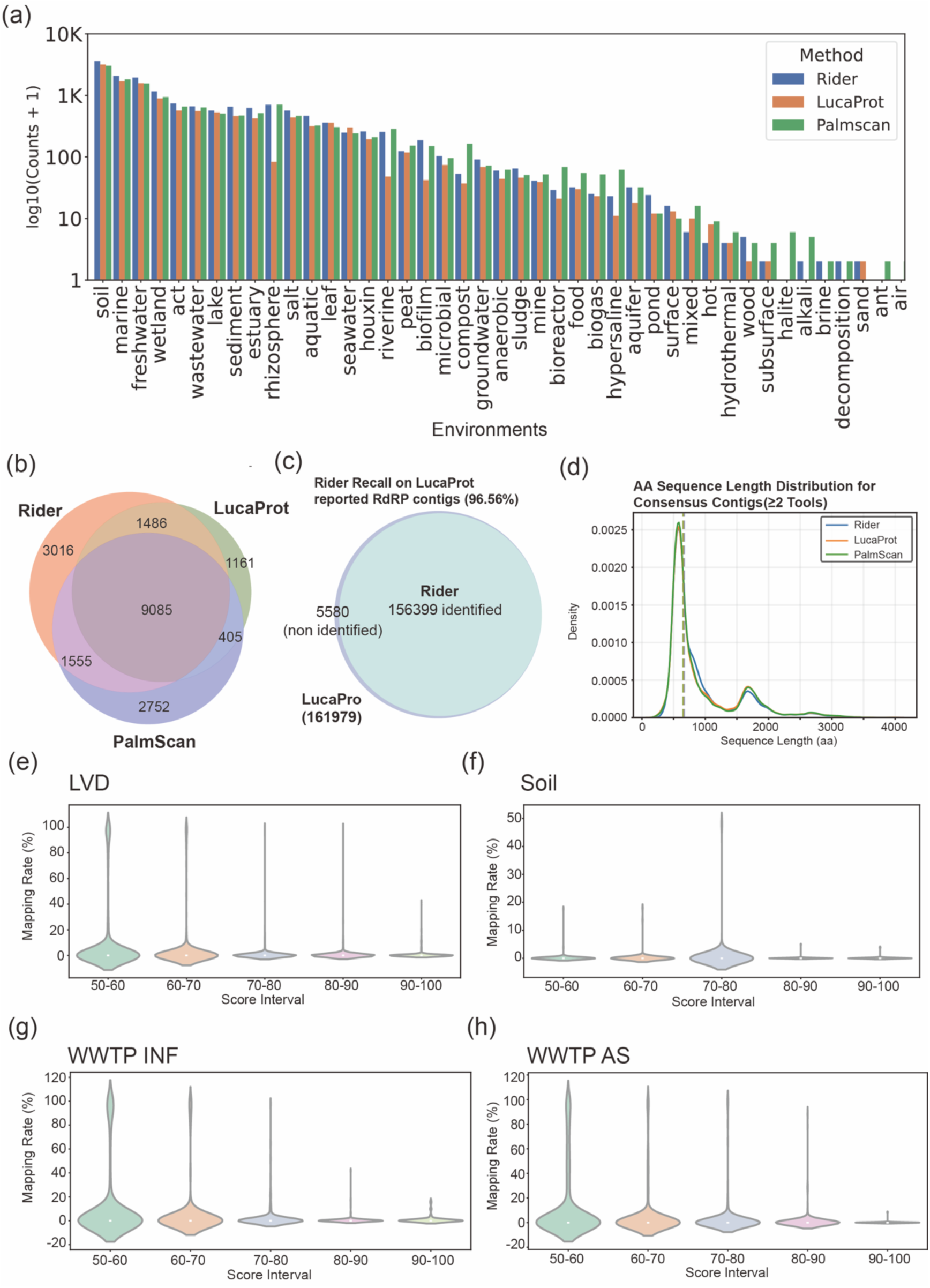
Rider unveils RNA viruses from diverse ecological systems with high confidance. **(a)** Distribution of RdRp-containing sequences identified by Rider (blue), LucaProt (orange), and PalmScan (green) across environmental biomes. Counts are log-transformed (log₁₀(count + 1)); soil, marine, and freshwater environments show the highest viral diversity. **(b)** Venn diagram showing overlap in RdRp sequence detection among the three methods. Rider uniquely identifies 8,384 sequences (>200 amino acids) with high structural similarity to known RNA viral proteins. **(c)** Rider achieves 95.81% recall of RdRp sequences reported by LucaProt, demonstrating high sensitivity on known viral proteins. **(d)** Amino acid length distribution of shared identified viral contigs by Rider and other tools. Rider detects a higher proportion of short sequences, consistent with improved performance on fragmented assemblies. **(e–h)** DNA read mapping of Rider-predicted viral contigs across four representative environments. Violin plots show mapping rates for contigs binned by Rider score. Contigs with high scores (90–100) consistently show near-zero DNA support, suggesting RNA origin and excluding DNA virus contamination. Environments shown include (e) LucaProt validate metatranscriptomes data (LVD) ^13^ and our sequencing data, including (f) soil, (g) wastewater influent (WWTP INF), and (h) activated sludge (WWTP AS).

Notably, DNA validation of Rider-predicted viral candidates from human-associated samples, such as those from the gut microbiome, was not included in this analysis due to the unique challenges associated with host-dominated metagenomes and low viral signal.

### Rider expands the RNA virus sequence space across diverse environments

To systematically assess Rider’s ability to discover RNA viruses from diverse ecological niches, we applied it to a large-scale environmental dataset comprising over 10,000 metatranscriptomic samples^13^. Predicted ORFs were clustered at 90% amino acid similarity, and representative centroid sequences were analyzed. Rider identified a total of 15,142 unique clusters of viral-like open reading frames (ORFs), each represented by a centroid sequence and comprising at least four member sequences (≥90% amino acid identity). These clusters were derived from Rider-predicted ORFs ≥200 amino acids in length, all treated as centroids for subsequent analyses.

Among these, Rider identified 12,126 contigs that were co-identified by at least one additional tool (LucaProt or PalmScan), providing orthogonal support for their viral origin. Collectively, these contigs encode 159,884 Rider-predicted ORFs, substantially expanding the known RNA viral protein space (Fig. 4a,b). Compared to LucaProt ^13^ and PalmScan^1,2^, Rider recovered 22.7% and 12.4% more viral-like ORFs, respectively, across 45 ecosystem types, with the highest richness observed in soil, marine, and freshwater samples (Fig. 4a, Fig. S6). For example, soil is the top RNA viral-abundant ecosystem, and Rider performed better in virus retrieval by 19.41% and 14.56% compared to PalmScan and LucaProt. These results underscore Rider’s broad sensitivity and capacity to uncover diverse RNA viruses from complex environmental data.

To further validate Rider’s ability to detect RdRp, we compared its predictions with those from LucaProt and PalmScan, both of which are RdRp-centric tools. A total of 9,085 contigs were co-identified by all three methods, and at least two tools supported 12,531 contigs. Among these, Rider contributed 12,126 contigs, more than LucaProt (n = 10,976) and PalmScan (n = 11,045), confirming its superior sensitivity in detecting RdRp-associated sequences. In addition to consensus predictions, Rider uniquely identified 3,016 proteins that exhibited significant structural similarity to known RNA viral proteins, as validated by Foldseek against the Rider structural reference database (Fig. 4b). The protein length distribution of predicted RdRp-like sequences showed strong concordance between Rider, LucaProt, and PalmScan, reinforcing Rider’s ability to accurately recover replication-associated proteins and its alignment with existing motif-based tools (Fig. 4d, Fig. S5).

To further benchmark its sensitivity, we evaluated Rider on a published environmental RNA virus dataset enriched for RdRp sequences from the same sample source. ^19^, Rider successfully recalled 96.56% (156,399 out of 161,979) of known RdRp-containing contigs, confirming its high recall for canonical viral markers (Figure 4d). This capability is further supported by the amino acid length distribution of contigs co-identified by at least two tools (Fig. 4d), showing an identical distribution curve compared to other SOTA methods regarding the length of sequences of potential candidates (Table S4). These results highlighted Rider’s ability to extract informative sequence features even in fragmented or domain-incomplete proteins.

### Extending to human gut virome: Insights from an IBD cohort

Profiling the human gut RNA virome remains a major technical challenge, especially in the context of complex diseases such as inflammatory bowel disease (IBD). Although recent studies suggest that the gut RNA virome may influence host health, findings across cohorts have often been inconsistent. These discrepancies reflect broader limitations in virome research, including extreme sequence diversity, fragmented assemblies, and sparse representation in reference databases ^37^.

To address these challenges, we applied Rider to a well-characterized IBD metatranscriptomic cohort comprising Crohn’s disease (CD), ulcerative colitis (UC), and non-IBD samples (Table S5-S8). Rider’s predictions were compared to those from a state-of-the-art motif-based method. Both tools revealed consistent trends in overall RNA viral richness and abundance, with CD samples exhibiting higher RNA viral levels than UC (p < 0.05; Figure 5a–c). Shared contigs (n = 104, identified by at least two methods) were further validated by structural alignment (Prob-score: 73.51, Figure 5j) and DNA reverse mapping (below 20% mapping rate, Figure 5j).

**Fig. 5.**
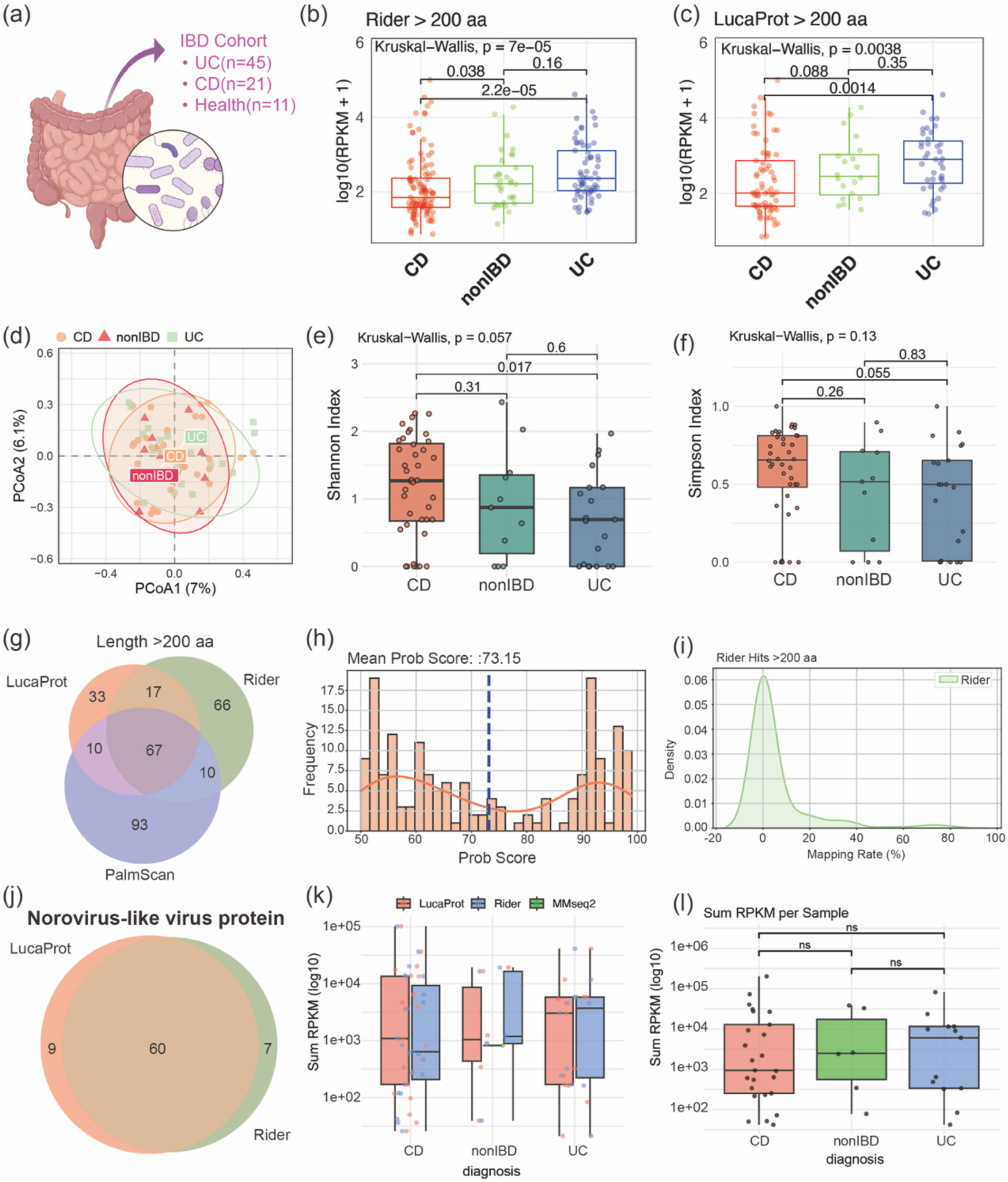
Rider enables robust profiling of the human gut RNA virome across inflammatory bowel disease (IBD) cohorts. **a**, Schematic overview of the IBD cohort, comprising individuals with ulcerative colitis (UC, n = 45), Crohn’s disease (CD, n = 21), and non-IBD healthy controls (n = 11). **b,c,** Relative abundance of predicted RNA viral contigs (log₁₀(RPKM+1)) identified by Rider (b) and LucaProt (c) across diagnostic groups, using a minimum protein length threshold of 200 amino acids. A total of 172 high-confidence candidates were retained for analysis after excluding sequences with >90% DNA read mapping rates. **d**, Principal coordinates analysis (PCoA) based on Bray–Curtis dissimilarity reveals beta diversity patterns of RNA virus profiles among diagnostic groups. **e,f,** Alpha diversity of RNA viral candidates measured using the Shannon (e) and Simpson (f) indices. No significant differences were observed between groups (Kruskal–Wallis test, P > 0.05). **g,** Venn diagram showing the overlap of contigs encoding viral-like proteins (>200 amino acids) predicted by Rider, LucaProt, and PalmScan, with annotations based on the Rider structural reference database (v1). **h,** Foldseek-based structural validation of Rider-predicted proteins yields a mean probabilistic Foldseek score (Prob-score) of 73.15, indicative of high-confidence structural similarity to known RNA virus proteins. **i,** Density distribution of DNA mapping rates for Rider-predicted candidates, below 20%, suggesting limited host contamination. **j,** Overlap of norovirus-like contigs predicted by Rider and LucaProt, with 60 sequences shared between methods. **k,l,** Group-wise comparisons of norovirus-like contig abundance (summed RPKM per sample) identified by Rider (k) and LucaProt (l), showing no statistically significant differences between diagnostic groups (Wilcoxon test, P > 0.05). The conventional sequence similarity method MMseq2 detected only a single norovirus-like sequence (see Supplementary Table 3).

To further explore the utility of Rider in characterizing disease-associated virome dynamics, we analyzed RNA viral patterns across diagnostic groups within the IBD cohort (Table S5). Relative abundance analysis (log₁₀(RPKM+1)) revealed consistent trends between Rider and LucaProt, with both tools showing significantly higher viral loads in UC samples compared to CD and non-IBD controls (Kruskal–Wallis P < 0.001; Fig. 5b,c). These findings suggest a potential enrichment of RNA viruses in UC-associated gut environments, independent of the detection method. Beta diversity analysis using Bray–Curtis dissimilarity (Fig. 5d) did not reveal clear separation between diagnostic groups, indicating that overall community composition remained broadly similar. However, alpha diversity metrics revealed more nuanced differences. Shannon and Simpson indices (Fig. 5e,f) both suggested that CD samples harbored the highest RNA viral diversity, while UC samples consistently exhibited the lowest. For Rider, the global difference approached significance (Kruskal–Wallis *P* = 0.057 for Shannon), and the pairwise comparison between CD and UC was statistically significant (*P* = 0.017, Wilcoxon test). In contrast, LucaProt showed a similar trend but did not reach significance in post hoc testing, highlighting Rider’s increased sensitivity in resolving subtle group-wise differences in viral diversity, probably owing to its superior sensitivity on shorter fragmented sequences.

Shorter viral signals were also important but usually overlooked roles in virome profiling study. Focusing on short viral signals represents an emerging direction in virome research. For instance, recent k-mer-based approaches have detected hundreds of short sequences associated with IBD ^38^, many of which lack canonical domain structures. We identified 156 RdRp and non-RdRp candidates with lengths ranging from 150 to 200 aa, with the highest probability score of 86 (Figure S9). This demonstrates that Rider’s ability to recover fragmented, short, and endogenous viral elements provides a more complete and unbiased view of the gut virome. Failure to detect such sequences may lead to systematic underestimation of viral abundance and diversity and may mask biologically relevant virus–host associations^39^. However, slightly increased contamination was observed via DNA mapping (Figure S9c); therefore, we need to treat these short fragments cautiously. The effect likely stems from the gut’s distinctive genomic complexity, such as horizontal gene transfer, the prevalence of long/short interspersed nuclear elements (SINEs/LINEs), and uncharacterized viral remnants. ^40^

### Detection of norovirus-like RNA viruses across IBD groups

Noroviruses are a major group of positive-sense, single-stranded RNA viruses in the family *Caliciviridae*, and have been implicated in gastrointestinal infections and potentially in the pathogenesis or exacerbation of IBD. However, existing studies offer conflicting results regarding their association with IBD, with some cohorts reporting increased prevalence in patients, while others find no consistent enrichment. To investigate this further, we curated a dedicated norovirus protein reference database comprising capsid and non-structural proteins from RefSeq and applied a structure-based annotation strategy to Rider- and LucaProt-predicted viral ORFs. All predicted proteins were modeled using ESMFold and aligned against the norovirus reference database using Foldseek. This approach revealed a highly consistent set of 60 norovirus-like ORFs that were co-identified across both methods (Table S7), supporting the robustness of these predictions. Quantitative comparison showed that abundance estimates for these contigs were nearly identical between Rider and LucaProt (Fig. 5k), and group-wise comparisons of summed RPKM values revealed no statistically significant differences across diagnostic groups (Kruskal–Wallis *P* > 0.05; Fig. 5l), suggesting that norovirus-like RNA viruses are broadly distributed in the human gut virome but may not be differentially enriched in IBD.

Notably, the conventional sequence similarity-based method MMseqs2 identified only a single norovirus-like candidate, underscoring the limited sensitivity of traditional approaches in detecting highly divergent or fragmented viral sequences within complex metatranscriptomes. These results emphasize the value of structure-guided annotation for functionally characterizing RNA viruses that would otherwise remain hidden using standard alignment-based methods. More broadly, they highlight Rider’s potential as a scalable and robust tool for medical microbiome studies, particularly in resolving the diversity and dynamics of the human RNA virome in health and disease.

## Discussion

This work demonstrates that Rider, a paradigm framework combining a fine-tuned large protein language model and leveraging protein structure alignment, can boost the identification of both known and novel RNA viruses. The promising ability for filtering initial viral candidates at the first step (sequence-level prediction), followed by fast structure alignment validation, makes it possible for global-sample-level application in real-world metatranscriptomic sequencing data at moderate and much less resource cost than SOTA tools.

We attribute Rider’s sensitivity to a deliberate, data-centric training strategy designed to overcome the limitations of protein language models, particularly their fixed-length receptive fields. Models like ESM2 process only the first 1,024 amino acids of an input, which can lead to truncation bias when important domains, such as RdRp, may fall outside the embedding window in long polyproteins or fragmented metatranscriptomic contigs. To address this, Rider’s training corpus was specifically designed to include both short RdRp fragments and long polyproteins containing RdRp alongside adjacent replication-associated domains (e.g., helicases, proteases, capsid/nucleocapsid elements, membrane anchors). This design enables the model to recognize not only canonical viral motifs but also contextual features from neighboring regions. As a result, Rider can correctly classify contigs that lack complete RdRp motifs, such as ORF1a-only sequences, by leveraging upstream or adjacent replication signals that motif-centric or alignment-based methods often miss. This ability to generalize beyond strict motif boundaries likely underpins Rider’s enhanced sensitivity to fragmented or non-canonical viral sequences, as reflected in the length distribution of identified ORFs (Figure 4e) and its superior performance on non-RdRp proteins (Figure S3).

The identification of RNA viruses from environmental and human metatranscriptomic assemblies remains fundamentally challenging due to the fragmented nature of contigs and the diversity of viral sequences, which often lack complete functional domains or conserved motifs such as those in RdRp. These limitations hinder traditional alignment-based methods (e.g., BLAST) that rely on high sequence similarity. Our results show that Rider consistently identifies a core set of viral contigs shared with SOTA tools, demonstrating its robustness in environmental RNA virome studies. Furthermore, Rider’s concordance with LucaProt in the IBD cohort with faster speed and similar results compared to LucaProt, highlighting its applicability in medical microbiome research and its potential to uncover viral sequences beyond the reach of classical motif- or alignment-based approaches.

While Rider shows strong sensitivity for both canonical and divergent RNA virus proteins, several limitations merit note. First, because Rider is trained to detect a broad spectrum of replication-associated signals (including regions upstream or adjacent to RdRp), its positives can include contigs that lack an explicit RdRp domain (Figure 3c,d, S10), thus can overcome the fragmented-in-feature assembly data in metatrascriptomic anslysis. For users who require strict RdRp retrieval, we recommend post-filtering Rider outputs with motif-based tools such as LucaProt, PalmScan (as shown in Figure 4b), or a curated structural RdRp-specific reference set to improve specificity; conversely, retaining raw Rider outputs is appropriate when the goal is maximal recovery of fragmented or non-canonical replication proteins. Second, structure validation relies on the breadth of the structure database, as shown by the decrease hitting rate of Rider on previously reported RdRp sequences (Figure 5g,j). Besides, systematic functional annotation and taxonomic assignment via structure alignment remain challenging. Structural prediction and alignment are useful supporting evidence but are not yet sufficient for definitive classification: current structure databases are incomplete, and fold similarity without genomic, syntenic, or ecological context can be ambiguous. Therefore, structure-based matches should be treated as indicative rather than conclusive.

Looking forward, we suggest three priorities to strengthen structure-aware viral discovery: (i) build benchmark datasets and standardized evaluation protocols for structure-guided annotation; (ii) expand and curate structural reference libraries for viral domains (especially divergent folds); and (iii) integrate contextual metadata (host, environment, genome organization) with fast structure-alignment and long-window homology tools. Practically, we advocate a hybrid workflow: use Rider for high-sensitivity scanning of metatranscriptomic assemblies, then apply targeted motif filters, long-window models, alignments, and structural analyses to refine domain annotations and taxonomic assignment.

## Acknowledgements

This work was supported by the Zhejiang Provincial Natural Science Foundation of China (LR22D010001) and the Research Center for Industries of the Future (WU2022C030). F.J. also acknowledges support from the Westlake University-Muyuan Joint Research Institute (Grant No 206000006022404) and the HRHI program (202309010) of Westlake Laboratory of Life Sciences and Biomedicine.

We thank Dr. Fa-Jie Yuan and Prof. Ren Sun, both from Westlake University, for their valuable suggestions and insightful discussions that helped improve this work.

